# Single Cell Profiling of CD45^+^ Spinal Cord Cells Reveals Microglial and B Cell Heterogeneity and Crosstalk Following Spinal Cord Injury

**DOI:** 10.1101/2022.03.29.486287

**Authors:** Elizabeth S. Fisher, Matthew Amarante, Natasha Lowry, Steven Lotz, Farhad Farjood, Sally Temple, Caitlin E. Hill, Thomas R. Kiehl

**Affiliations:** Neural Stem Cell Institute, Rensselaer, NY 12214

## Abstract

It is well established that immune cells play crucial roles after spinal cord injury (SCI). However, our knowledge of the contributions of various immune cells to injury progression and repair is incomplete. These gaps in understanding hamper development of SCI therapeutics. In the current study, using single-cell RNA sequencing, and transcriptomic analysis, the populations of resident and circulating CD45^+^ immune cells present within the uninjured and injured mouse spinal cord were identified. In the uninjured and subacutely-injured (7 day) spinal cord, most CD45^+^ cells were microglia while in chronic SCI (60 day) B cells predominated. Examination of microglia and B cell clusters showed subtype-specific alterations after SCI, including the presence of both immature and mature B cells chronically. Analysis of the expression of signaling partners in B cells and microglia identified injury-related microglia-B-cell interactions. This sequencing resource establishes unidentified interactions revealing new mechanisms to target inflammatory responses for SCI repair.

## Introduction

Spinal cord injury (SCI) initiates complex molecular cascades signaling both tissue destruction and remodeling^1, 2^. The cellular and molecular mechanisms leading to persistent deficits after SCI are only partially understood. Recently, profiling of individual cells within the spinal cord and their temporal responses after SCI by single-cell RNA sequencing (scRNA-seq) has substantially advanced our understanding of the breadth of cellular plasticity following SCI, and revealed concurrent dynamics within specific cell types including neurons, glia, and immune cells^3–6^. A recent scRNA-seq study of immune cells in severe SCI showed prolonged gene expression changes in microglia (up to three months post injury) which were proposed to enhance recovery^4^. The interactions between different immune cell classes, especially in chronic injury, remain largely unknown.

The immune response is a key mediator of tissue damage and wound healing. It intricately coordinates cellular responses of both the innate (e.g., neutrophils, macrophages) and adaptive (e.g., T-cells, B cells) immune systems. After SCI, microglia, the CNS resident immune cells, and infiltrating immune cells become activated and initiate diverse and complicated damage responses^7^. Prior studies have examined individual immune cell classes and their responses following SCI^8–13^ including neutrophils^14–16^, macrophages^17, 18^, T cells^19, 20^, and B cells^21^. SCI fails to elicit coordinated immune cell responses necessary for effective tissue repair^22^ leading to chronic inflammation, poor wound resolution, and tissue fibrosis^2, 3, 18, 23–25^. To effectively develop therapeutic interventions to mitigate SCI tissue damage, prevent chronic inflammation, and initiate effective remodeling, a more complete temporal understanding of the immune response and interactions occurring between infiltrating and resident immune cells is required.

In the current study, we profiled resident and infiltrating CD45^+^ cells in the intact and injured spinal cord at subacute and chronic stages using scRNA-seq to establish how the various immune cells change and respond to the evolving injury. CellPhoneDB was used to probe for specific cell-cell interactions warranting further exploration as targets for SCI repair. Based on the gene expression profile of individual immune cells, we show that SCI elicits pronounced changes in microglia and B cell numbers and functions. Microglia undergo SCI-specific changes distinct from those occurring following brain injury, with a population appearing acutely and persisting chronically. SCI causes formation of ectopic lymphoid follicles at the injury epicenter containing immature and maturing B cells. The microglial and B cell responses arising following injury and persisting chronically include the expression of multiple chemokine receptor-ligand pairs indicating cellular crosstalk. Together, our results reveal that SCI induces chronic modifications in both resident and infiltrating immune cells. This resources identifies persistent cell-cell interactions between and within B cells and microglia that can be mined to address their role in pathobiology and chronic degeneration seen in the cord following injury.

## Results

### Single-cell sequencing of CD45^+^ cells after spinal cord injury

To establish the cellular and transcriptional profile of resident and infiltrating immune cells in the spinal cord, adult (10–12-week-old) female Swiss-Webster mice received a moderate thoracic spinal cord contusion injury (T9, 6.25 mm, NYU device). Three mm sections surrounding the lesion epicenter were pooled, dissociated, and processed from injured versus uninjured control mice (n=6 mice/condition) as summarized in **Figure 1a-c**. Individual immune cells were isolated by FACS using the pan-immune cell marker CD45, and scRNA-seq was performed using the nanowell-based iCell8 system (Takara Bio USA), which generates more reads per cell than a droplet-based system. In total, 426 cells from uninjured, 102 cells from 3 dpi, 454 cells from 7 dpi, and 945 cells from 60 dpi passed quality control and were analyzed (**Figure 1c**). The average total transcript reads across conditions was 645,000 reads per cell. Sample reads at each time point were well within the range necessary for downstream batch correction and analyses to robustly discriminate different cell populations^26^. Following SCTransform^27^ to normalize the data and remove technical artifacts, samples were plotted by time to demonstrate proper normalization (**Supplemental Figure 1**). Fewer cells and substantially fewer reads (median of 5,423 reads/cell) were detected in the 3-dpi sample compared to the others. While this did not affect the analysis in context of the whole data set, disparate read counts and sparsity from this sample introduced excess variability into downstream analysis of smaller subsets. As such, after the initial clustering analysis (**Figure 2**, **Supplemental Figure 2**), the 3 dpi data was excluded from subsequent analyses.

**Figure 1.**
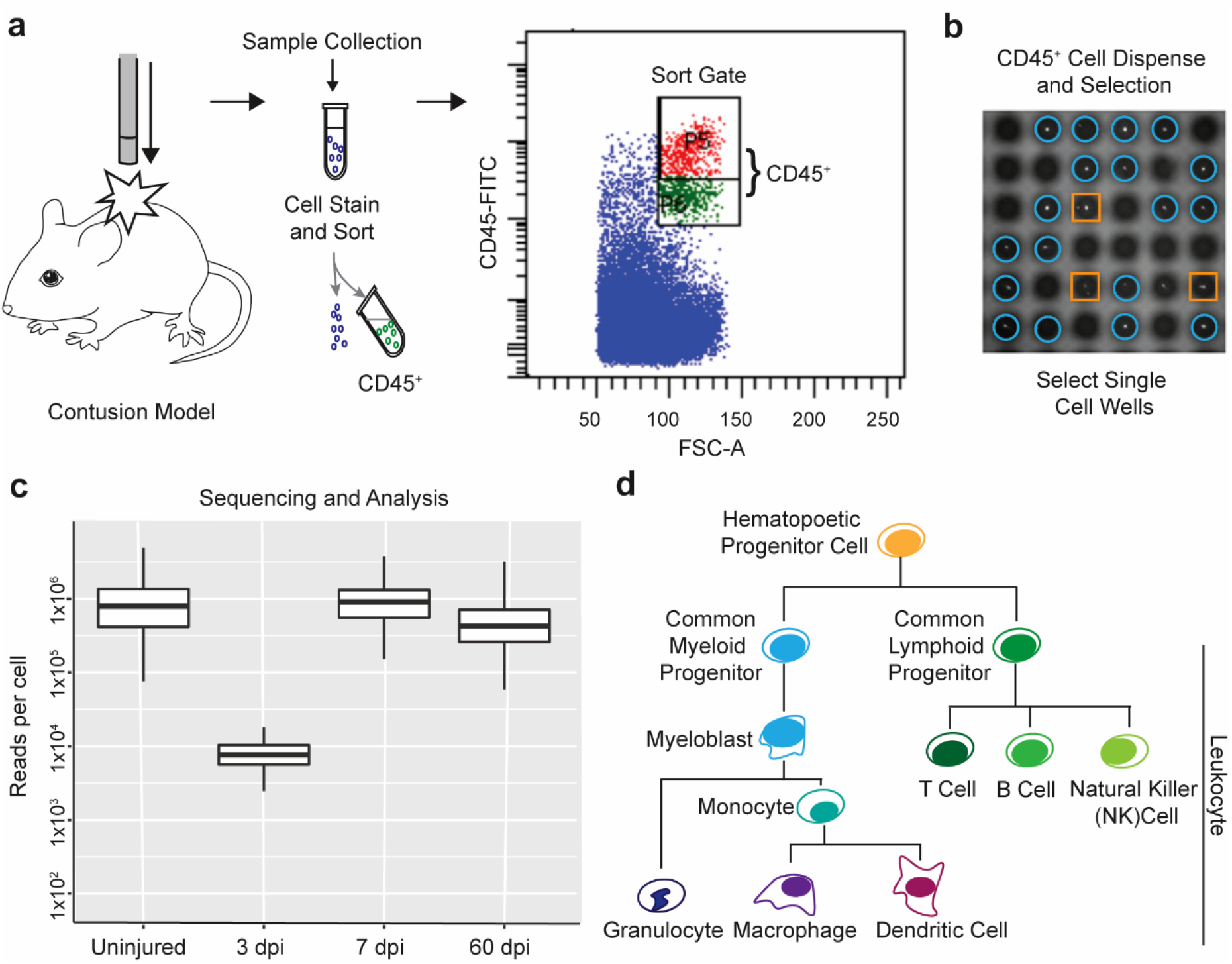
Workflow and quality control for single-cell isolation and sequencing of spinal cord immune cells. a) Schematic of experimental approach. b) Isolation of single cells onto the iCell8 chip. Wells containing single live cells (blue circles) were identified by microscopy imaging for further processing; wells with multiple cells or dead cells (orange squares) were excluded. c) Reads per cell for the four conditions, represented by a box and whiskers plot. Box represents median and quartiles, while the whiskers are maximum and minimum values, excluding outliers. d) Schematic of the immune cell lineage, simplified to include cell populations relevant to this study.

**Figure 2.**
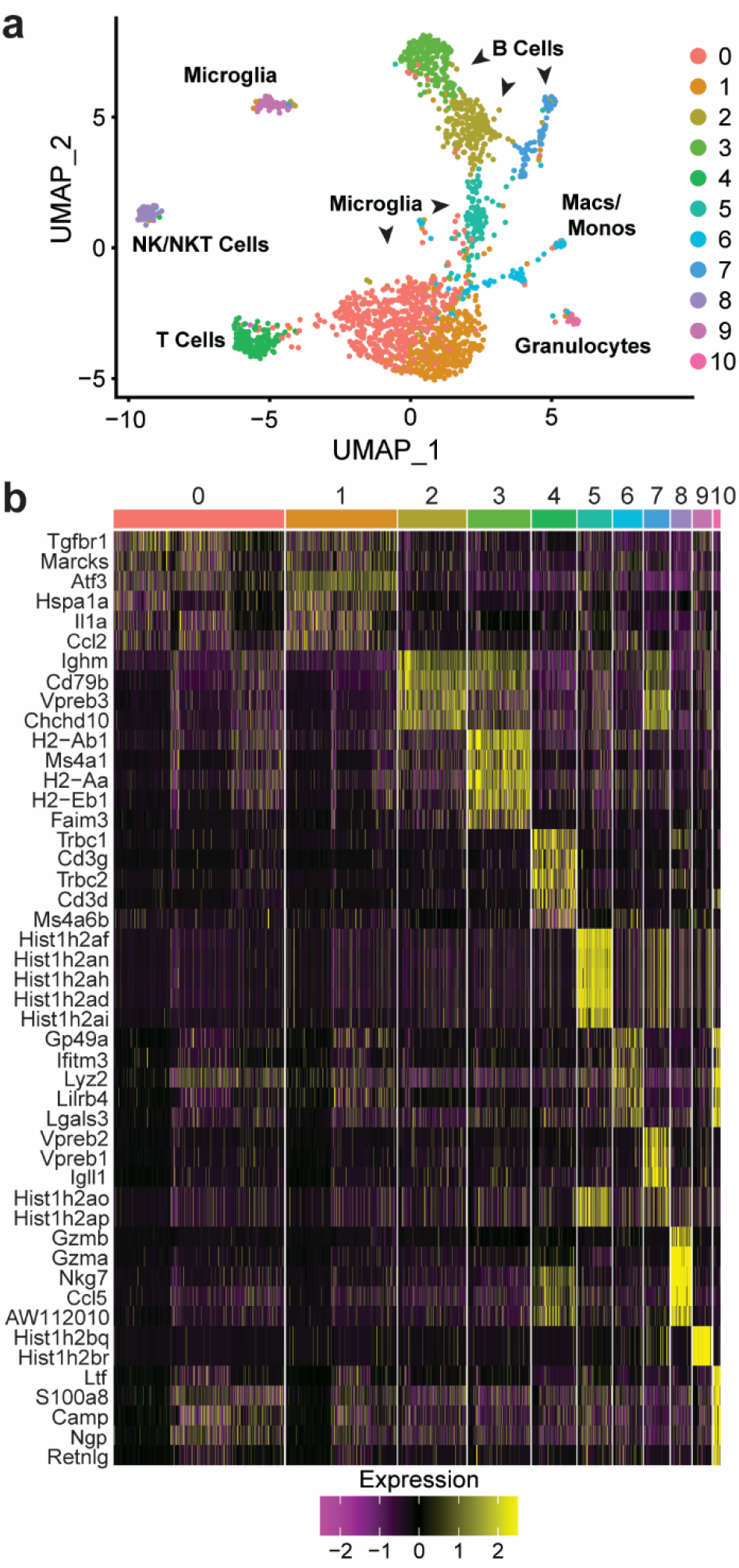
Analysis of combined data across all timepoints reveals clusters containing major immune cell types. a) UMAP embedding of all timepoints combined after normalization by SCTransform. Clusters clearly separated into distinct immune cell populations. Cluster 0, 1, 5 and 9 express canonical microglia markers. Cluster 2, 3 and 7 express B cell markers. Cluster 4 expresses markers of T cells, and cluster 8 expresses NK cell markers. Cluster 6 expresses markers of monocytes/macrophages, and cluster 10 expresses granulocyte markers. b) Heatmap of the top 5 genes driving cluster separation.

### Characterization of immune cell diversity after SCI

Samples from all time points were combined and clustered together using the shared nearest neighbor algorithm to gain an overview of the complete immune cell dataset. Eleven cell clusters (Clusters 0–10) were identified and subsequently mapped to specific immune cell types (UMAP: **Figure 2a**). The identified cell clusters included the major innate and adaptive immune cell types known to participate in the response to SCI^23, 28^ including microglia (clusters 0, 1, 5, and 9), NK/NKT cells (cluster 8), macrophages/monocytes (cluster 6), granulocytes (cluster 10), T cells (cluster 4) and B cells (clusters 2, 3, and 7) (**Figure 2a**). The top 5 DE genes for each cluster are presented as a heatmap (**Figure 2b**). Cell type designations were first established by analyzing differentially expressed (DE) genes in each cluster and manually comparing them to several canonical markers of leukocytes and microglia. Cell type designations were confirmed using both SingleR^29^, a nearest neighbor, reference-based, label-transfer approach based on Spearman correlations, and by comparing the results with the cell-type reference ImmGen database (**Supplemental Figure 2a,b**).

### Dynamic temporal changes in immune cell types occur after SCI

Microglia (Cluster 0, 1, 5 and 9) and B cells (2, 3 and 7) comprised the bulk of the total CD45^+^ cells combined across timepoints (**Figure 3a, b**) (53.8% and 28.5% of the total cells, respectively). The remaining 17.7% of total cells were macrophage/monocytes (5.2%), NK/NKT cells (3.5%), neutrophils (1.3%) and T cells (7.7%).

**Figure 3.**
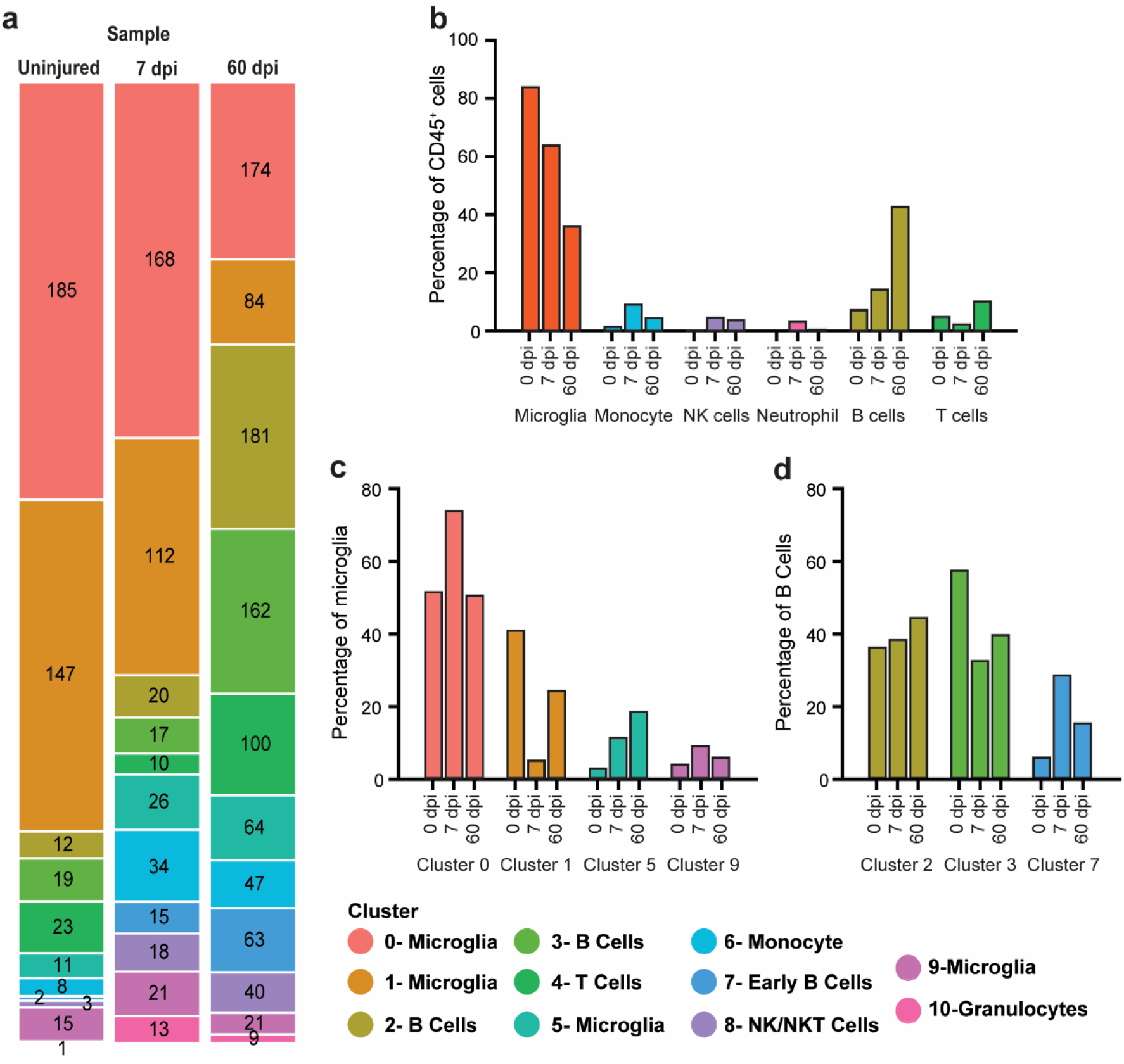
Immune cell populations and the sub-types of microglia and B cells present within the spinal cord change over time in response to SCI. a) Stacked bar chart showing the number of cells present within each cluster at each time point as a proportion of all CD45^+^ cells isolated. b) Bar chart of the percentage of major immune cell types present in the spinal cord and how they change in response to injury. c) and d) bar charts depicting the percentage of microglia (c) and B cells (d) in each cluster relative to all microglia or B cells, respectively. Note the change in percentage of the different clusters within each cell type in response to injury.

The profile of the immune cells within the spinal cord changed in response to injury progression. In the uninjured spinal cord, the vast majority of CD45^+^ cells isolated were microglia (84%). The next largest cell population was B cells (7.7%). Peripheral myeloid, NK, and T cells made up the remainder (8.3%). After SCI, microglia predominated subacutely (64.1% at 7 dpi); but, in chronic SCI, they comprised just 36.3% of total CD45^+^ cells. In contrast, B cells that were initially 7.7% of total CD45^+^ cells in the uninjured tissue increased to 14.7% at 7 dpi and became the largest population (43%) in the chronic state. Other cells identified showed smaller changes in response to injury (**Figure 3a, b**). T cells decreased from ∼5% of CD45^+^ cells in the uninjured cord to 2.8% at 7 dpi, rebounding to 10.6% of cells isolated chronically. Peripheral myeloid and NK cells were present at all time points post-SCI, but their numbers were low, ranging from 2-10% for monocytes/macrophages, 0-4% for granulocytes, and 1-5% for NK cells. Collectively, this analysis revealed substantial alterations in immune cell populations present within the acutely and chronically injured spinal cord, particularly in microglia and B cells.

Microglia and B cells were distributed across multiple clusters, implicating different cell activation states or subtypes^30^. Analysis of the proportion of microglia (**Figure 3c**) and B cells (**Figure 3d**) by their respective clusters showed time-specific changes warranting further analysis.

### Uninjured spinal cord microglia are primed to respond to injury

SCI induces robust inflammation and greater scar formation than a similar magnitude brain injury^31, 32^. Microglia are hypothesized to be involved, as microglia-astrocyte interactions drive glial scar formation^33, 34^. To identify regional-specific functional differences between brain and spinal cord microglia in inflammatory responses, we compared our uninjured spinal cord microglial transcriptomes with previously published brain microglia transcriptomes^35^. This identified several distinct gene expression profiles and GO pathway enrichments for microglia in each tissue (**Figure 4a**, **Supplemental Table 1**). Brain microglia were enriched for functions related to RNA processing, including translation, protein synthesis and protein processing. In contrast, spinal cord microglia were enriched for processes involved in peripheral immune cell recruitment and oxidative stress related processes, such as "PERK-mediated unfolded protein response" and nitric oxide biosynthetic and metabolic processes. This analysis demonstrated in the normal CNS, functions of brain and spinal cord microglia differ. Based on the endogenously expressed resident microglia genes, spinal cord microglia appeared primed to respond to injury. These differences could make them more responsive to injury, contributing to the greater inflammatory response after SCI than seen after brain injury^31^.

**Figure 4.**
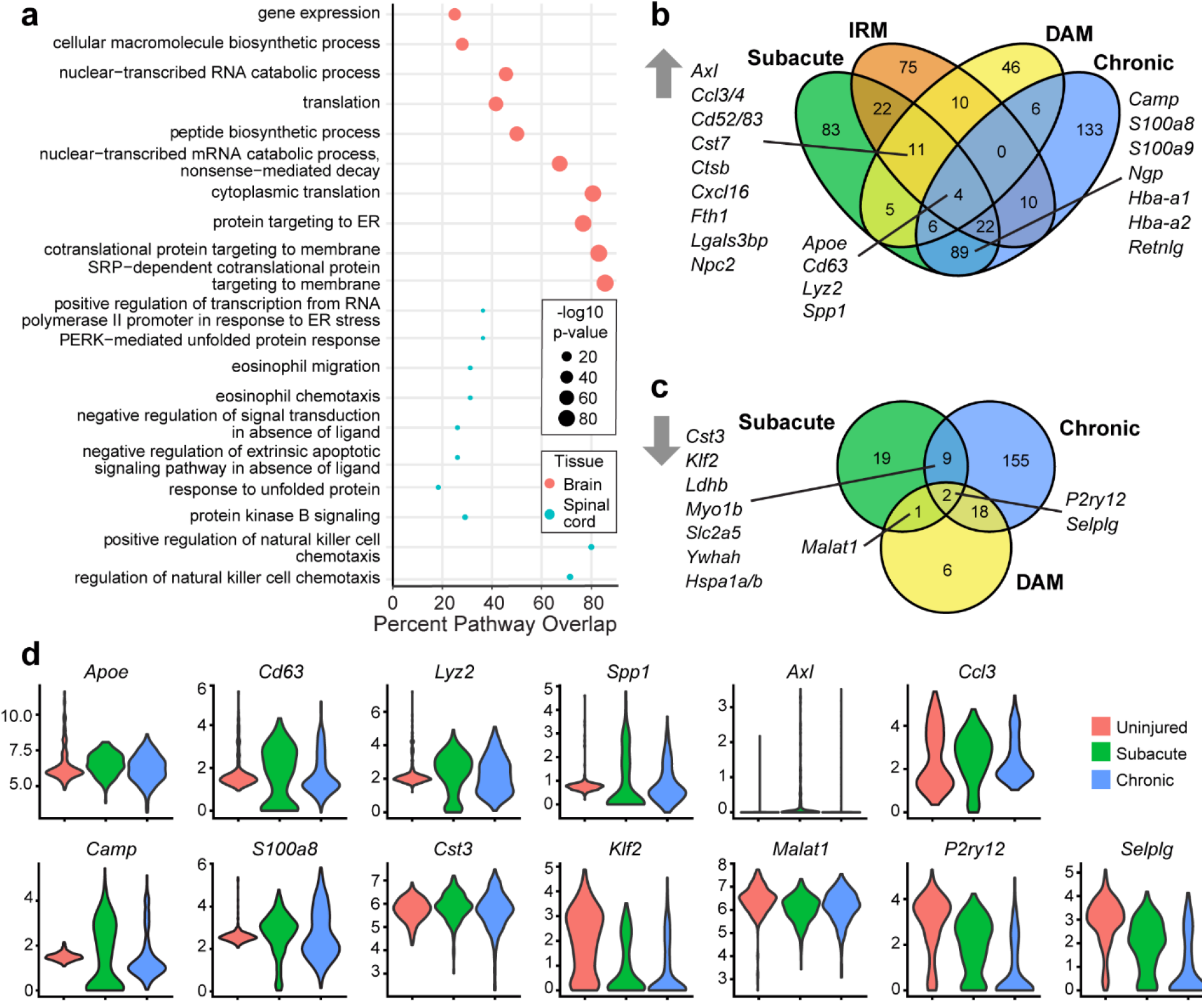
Comparison of uninjured, subacute, and chronic SCI microglia to uninjured and injured brain microglia. a) GO pathway enrichment in uninjured spinal cord and brain microglia^35^ using DE genes, represented as a dot plot (See **Supplemental Table 1** for additional enrichments). The x axis is the percentage of genes represented in the pathway which overlapped with enriched genes. b) Venn diagram of genes significantly upregulated in the subacute and chronic phase of SCI versus Injury Response Microglia (IRM^35^) and Disease-Associated Microglia (DAM)^36^ (See **Supplemental Table 2** for lists of up and down regulated genes following SCI.) c) Venn diagram of genes significantly downregulated in subacute and chronic SCI versus DAM. d) Violin plots of select significantly altered genes overlapping across injury states in our SCI data.

### Spinal cord and brain microglia respond differently to injury

Similarities between spinal cord microglia and disease associated microglia (DAM)^4^ have recently been identified. Activated brain and spinal cord microglia were compared using up- (**Figure 4b**) and down- (**Figure 4c**) regulated microglial genes in subacute and chronic SCI to those previously identified in brain injury and disease. For brain microglia, we used signature genes previously established for DAM from Alzheimer’s Disease (AD) mouse brains^36^, and injury responsive microglia (IRM^35^) generated by injection of lysolecithin into brain white matter. For down-regulated genes, the comparison was restricted to SCI-microglia and DAM because downregulated IRM genes were not available for analysis.

Tissue- and temporal-specific changes in gene expression were identified for spinal cord and brain microglia in response to injury (**Figure 4b, c**). Tissue specificity of the SCI-microglia response was highlighted by the identification of only six overlapping genes across all conditions. SCI-microglia shared four upregulated genes with DAM and IRM, *Apoe, Cd63*, *Lyz2*, and *Spp1* (**Figure 4d**), and two downregulated genes with DAM, the canonical microglia markers^35–37^ *P2ry12* and *Selplg* (**Figure 4d**). Temporal evaluation post-SCI identified eleven upregulated genes in acute SCI shared with DAMs and IRMs. These genes were related to metabolism (*Cst7*, *Ctsb*, *Npc2*, and *Fth1*a), and cell migration/adhesion (*Ccl3*, *Ccl4*, *Cxcl16*, *Cd63, Lgals3bp* and *Axl*). They also shared one downregulated genes with DAM, *Malat1*. A subset of the common genes is shown in **Figure 4d**. The DAM and IRM microglia genes overlapping with acute SCI included *Axl*, a gene upregulated in brain microglia in almost all disease states^38^. Chronic SCI-microglia had no upregulated genes in common with DAM and IRM, but eighteen downregulated genes were shared with DAM, including canonical microglia genes *Cx3cr1*, *P2ry13*, and *Csfr1*. This indicated that while some SCI-microglia responses resemble those in the injured brain^4^, most were unique to SCI. This could contribute to altered injury magnitudes between different CNS sites.

Comparison of SCI-microglia gene expression between acute and chronic showed temporal effects. Acute and chronic microglia shared some genes (89 upregulated and nine downregulated), but more genes were uniquely upregulated (83 and 133) or downregulated (19 and 155), under the respective conditions. Thus, not only did SCI-microglia differ from brain-microglia, but their gene expression profiles were specific to SCI injury phase. Multiple functions are ascribed to microglia^7, 39, 40^; understanding their context-specific changes and functional impact could aid in modulating their responses in injury and disease.

### Chronic response of B cells following SCI

The composition of B cells present after SCI, their functional profile, and the mechanisms leading to their accumulation are not yet known. The prevalence of B cells in our dataset — 25% of the total CD45^+^ cells across timepoints and 43% of cells isolated at 60 dpi — indicated roles for B cells in spinal cord pathophysiology beyond those currently described^21, 41^.

First, we examined mouse spinal cord sections by immunohistochemistry for B220^+^, a general marker of B cells. B220^+^ cells were found clustered in large discrete foci within the spinal cord at 42 dpi, (**Figure 5a-a’’**). Quantification of the number and area of clusters showed they formed between 3 and 28 dpi (**Figure 5b**) and increased in size between 28 and 42 dpi (**Figure 5c**, **Supplemental Figure 3a-b**). Similar accumulation of B cells with time post-SCI has been reported and hypothesized to contain of a mixed cell population^21, 41^.

**Figure 5.**
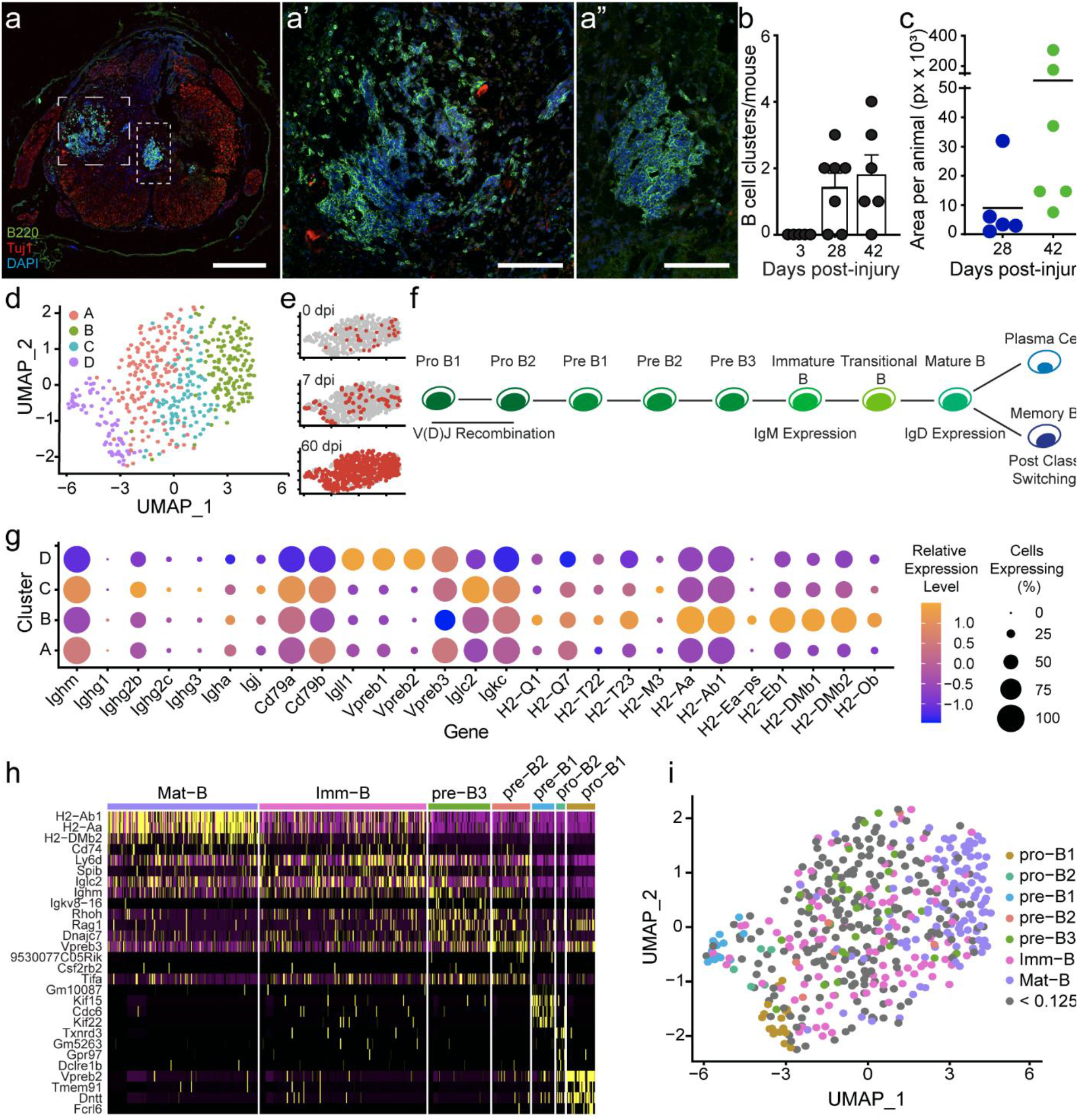
B cells increase in prevalence over time following SCI and contain multiple subtypes. a) Immunohistochemical staining of B cells in spinal cord tissue 42 dpi, stained with B220 (Green), Tuj1 (Red), and DAPI (Blue). Large and small dashed boxes indicate the regions shown in high power in a’ and a,” respectively. Scale bars: a) 500 μm, a’ and a”) 100 μm. b) Quantification of number of clusters identified in SCI animals over time following injury. Each dot represents an individual animal. Error bars represent SD. c) Quantification of the area of clusters per section, represented in pixels. d) Isolated and re-clustered UMAP of clusters 2, 3 and 7 identifies four subclusters of B cells. e) UMAP plot highlighting cells isolated from each timepoint, top is 0 dpi, middle is 7 dpi, and bottom is 60 dpi, demonstrating most B cells were isolated at 60 dpi. f) Dot plot demonstrating relative abundance and expression of the Ig class detected, of selected markers of early B cells, and selected MHC-I and MHC-II genes. g) Schematic showing the developmental lineage of B cells including important events along their development. H) Heatmap of reclustered data, and genes identifying them as a subtype identified by Jensen^42^. I) UMAP of B cell clusters, with cells identified by developmental stage.

To profile the B cells further, a secondary cluster analysis was performed using the three original clusters (Clusters 2, 3, 7). Four distinct sub-clusters, denoted A-D, were delineated (**Figure 5d; Supplemental Figure 4;** heatmap of top DE genes in the clusters) — including one, Cluster D, which emerged post-SCI (cf., **Figure 5d** and **5e**). B cell development involves progression from Pro-B1 cells through Mature B, which are activated to either produce antibodies (Plasma cells) or act as antigen presenting cells (APCs) (**Figure 5f**). To establish the activation and maturation states of cells present, several analyses were performed.

Expression of components of the B cell receptor (BCR) were examined to identify maturation state and whether any B cells were acting as APCs. Components analyzed included immunoglobulin heavy (Igh) and light chains (Igl, Igk), signaling components Ig-a (Cd79a) and Ig-b (Cd79b), Ig subclass type (i.e., IgM, IgD, IgG, IgA, IgE), and expression of MHC I and MHC II (**Figure 5g**). Cluster D, which appeared after SCI, had the most immature BCR. It contained cells expressing Igl components and enzymes needed for heavy chain V(D)J recombination, typically found in pro/pre B cells. Cluster A cells expressed intermediate levels of genes suggesting cells development of BCR expression. Cluster C cells expressed high levels of IgM and IgG2b indicative of activated B cells post-class switching. Cluster B cells expressed MHC-II components found in APCs. These results indicate the presence of B cells in multiple states of development and activation.

To identify their developmental state more precisely, our B cell data was compared to bulk RNA sequencing data from defined B cell linage states^42^ made into a reference in SingleR^29^. This analysis identified different B cell developmental stages from Pro-B1 cells through Mature B cells (**Figure 5h**). UMAP plotting of B cell identities showed Cluster D contained pro-B1, pro-B2, and pre-B1 cells, Cluster A was a mixture of pre-B2, pre-B3 and immature B cells, Cluster C included both immature and mature B cells and Cluster B was comprised of mature B cells (**Figure 5i**). Thus, a range of the B cell lineage existed within the spinal cord (**Figure 5h, i**), including a population of pro/pre-B cells emerging after SCI.

Localization of pre-B cells to the spinal cord meninges was recently reported^43^. Comparison with B cells isolated from the meninges of SCI and SOD1 mice^43^, a model of amyotrophic lateral sclerosis (ALS), showed the B cells we isolated from the spinal cord were most similar to those found in the meninges under inflammatory conditions (**Supplemental Figure 5a,b**). This demonstrated immature B cells are not restricted to the meninges after SCI. Understanding the functions of the B cells within the spinal cord and factors driving their entry could lead to new interventions to counteract B cell-mediated spinal cord pathology^21, 41^.

### Immune cell pathway responses after spinal cord injury

SCI induces complex immune-mediated changes^44^. Using our scRNA-seq data set, functional analysis of the GO biological pathways and semantic similarity measurements were used to identify shared or divergent pathways across time in response to injury (Top 10 terms, **Figure 6a,b;** complete dataset, **Supplemental Table 3**). We focused our analysis on the cell type specific functional enrichments occurring in response to SCI for microglia and B cells. The functions of other immune cells are included in **Supplemental Figure 6**. Top enrichments unique for each cell type and time point analyzed were shown, indicating specific, evolving roles of the various immune cells after SCI.

**Figure 6:**
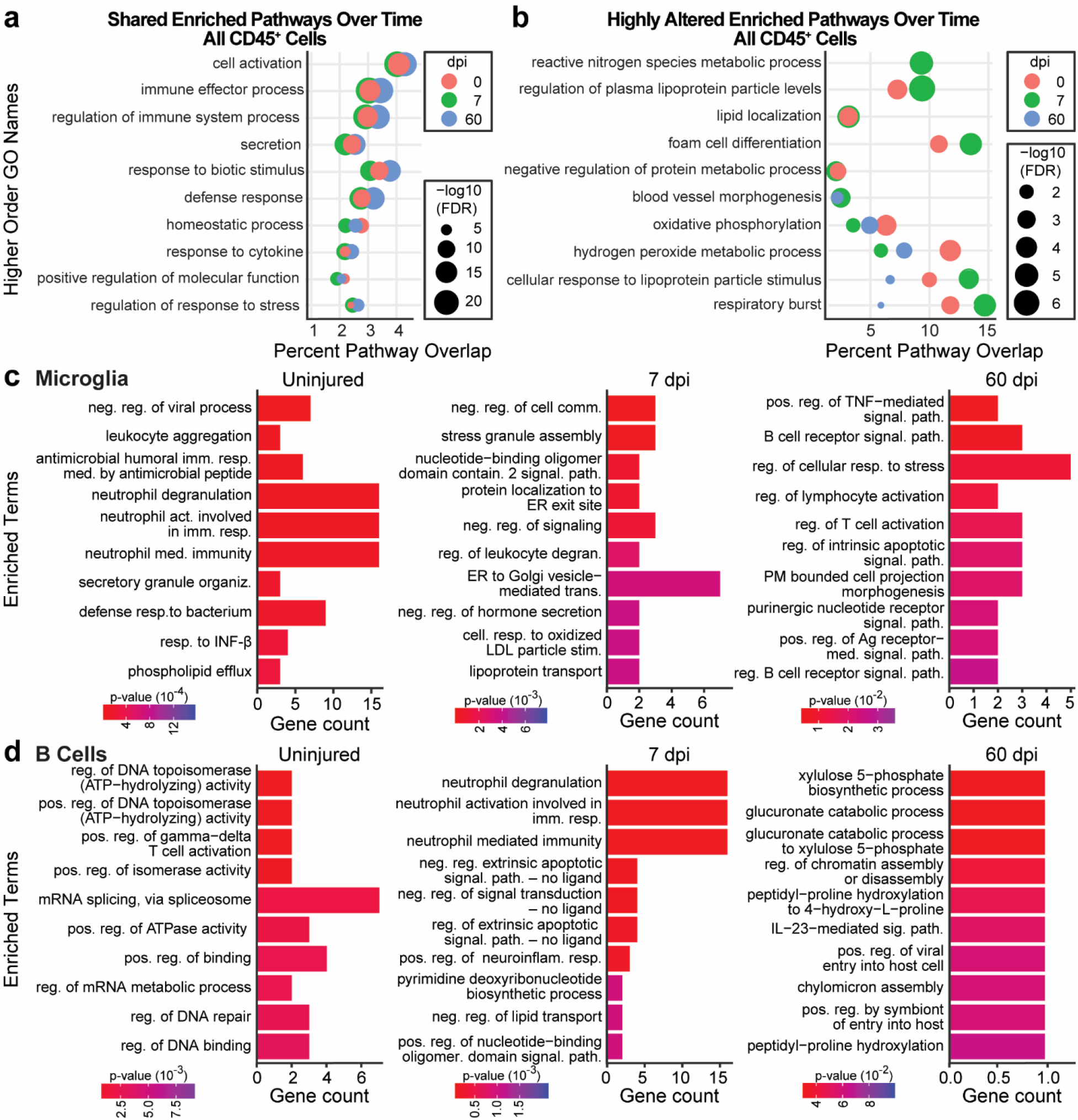
Identification of major pathways associated with timepoint and cell identity. a,b) Per timepoint, the Wilcoxon rank sum test was used on all cells isolated, followed by GO enrichment using the hypeR package. Semantic similarities of enriched GO terms were generated using rrvgo package, producing new categories displayed as a dot plot. The size of the dot indicates the False Discovery Rate (FDR) for the enriched category, and the x axis indicates the percentage of genes overlapping that pathway (See also **Supplemental Table 3** for enrichments). a) Enriched pathways with minimal change between timepoints. b) Enriched pathways either changed with time or were not represented at all timepoints. c,d) The Wilcoxon rank sum test was used for microglia (c), or B cells (d) at each timepoint, followed by GO enrichment using the Enrichr package and the “GO Biological Pathway 2021 library” to generate bar plots of significantly enriched pathways at each timepoint for the individual cell types (See **Supplemental Table 4** for additional enrichments).

Microglia are among the first cells to respond to injury, and have complex interactions with axons, glia and immune cells^24^. Our functional assessment showed microglia in the uninjured, subacute-SCI, and chronic-SCI spinal cord had distinctly different functional profiles related to changing interactions with other immune cells (**Figure 6c**). Uninjured microglia were enriched for terms reflective of a role in immune surveillance and interactions with neutrophils and other leukocytes; terms identified included “negative regulation of viral process,” “leukocyte aggregation,” and several associated with neutrophil biology. In subacute-SCI, microglia were enriched for pathways related to peripheral immune cell infiltration and stress responses including “regulation of leukocyte degranulation,” “negative regulation of cell communication,” “stress granule assembly,” and “lipoprotein transport.” In chronic-SCI, pathways linked to microglia-lymphocyte interactions predominated including “regulation of lymphocyte activation,” “B cell receptor signaling pathway,” “regulation of B cell receptor signaling pathway,” and “regulation of T cell activation.” Identification of terms in microglia related to their interactions with neutrophils and leukocytes aligns with their role as immune mediators between the CNS and periphery; currently, little is known about these interactions in the context of SCI. Detecting chronic microglia-lymphocyte interactions was particularly notable as it suggested a role for microglia in chronic B cell accumulation.

Unlike microglia, limited functions have been ascribed to B cells in the context of SCI^21, 41^. Our functional analysis of B cells (**Figure 6d**) showed diverse functions related to antibody refinement and interactions with other immune cells. Uninjured spinal cord B cells were enriched for functions related to DNA and RNA modifications suggestive of antibody refinement such as “regulation of DNA binding,” “regulation of DNA repair,” “regulation of DNA topoisomerase activity,” “positive regulation of isomerase activity,” and “mRNA splicing, via spliceosome.” Subacute SCI-B cells were enriched for multiple pathways regulating neutrophils, including “neutrophil degranulation,” “neutrophil activation involved in immune response,” and “neutrophil mediated immunity;” while chronic SCI-B cells were enriched for “interleukin-23 mediated signaling pathway” and several metabolism pathways. IL-23 signaling coordinates germinal center class-switching and promotes germinal B cell centers^45, 46^. Interestingly, both neutrophils and astrocytic-expression of IL-23 are linked to B cell accumulation and pathology in the experimental autoimmune encephalomyelitis (EAE) model of multiple sclerosis^46^.

### Interactions between microglia and B cells in the spinal cord

To identify potential cell-cell interactions between immune cells, we applied the publicly available repository of curated receptors, ligands and their interactions, CellPhoneDB^47^, to our dataset. As we were interested in identifying mechanisms contributing to the accumulation of B cells within the chronic SCI to test in future studies, and microglia were by far the majority of cells in our dataset, we focused on investigating microglia-B cells interactions; additional interactions are shown in **Supplemental Figure 7**.

In total, twenty-five ligand-receptor interactions were identified between B cells and microglia (**Figure 7a-d**). CellPhoneDB pairings are directional: eleven microglia-B cell (M–B) and 14 B cell–microglia (B–M) interactions were found. The number of M–B/B–M pairings decreased with injury progression: 14, 12, and 9, were identified for uninjured, 7 dpi, and 60 dpi spinal cords, respectively. Examination of the identified receptor-ligand pairings predict that a subset has roles in microglial survival, immune cell infiltration, and neuroprotection warranting further investigation.

**Figure 7.**
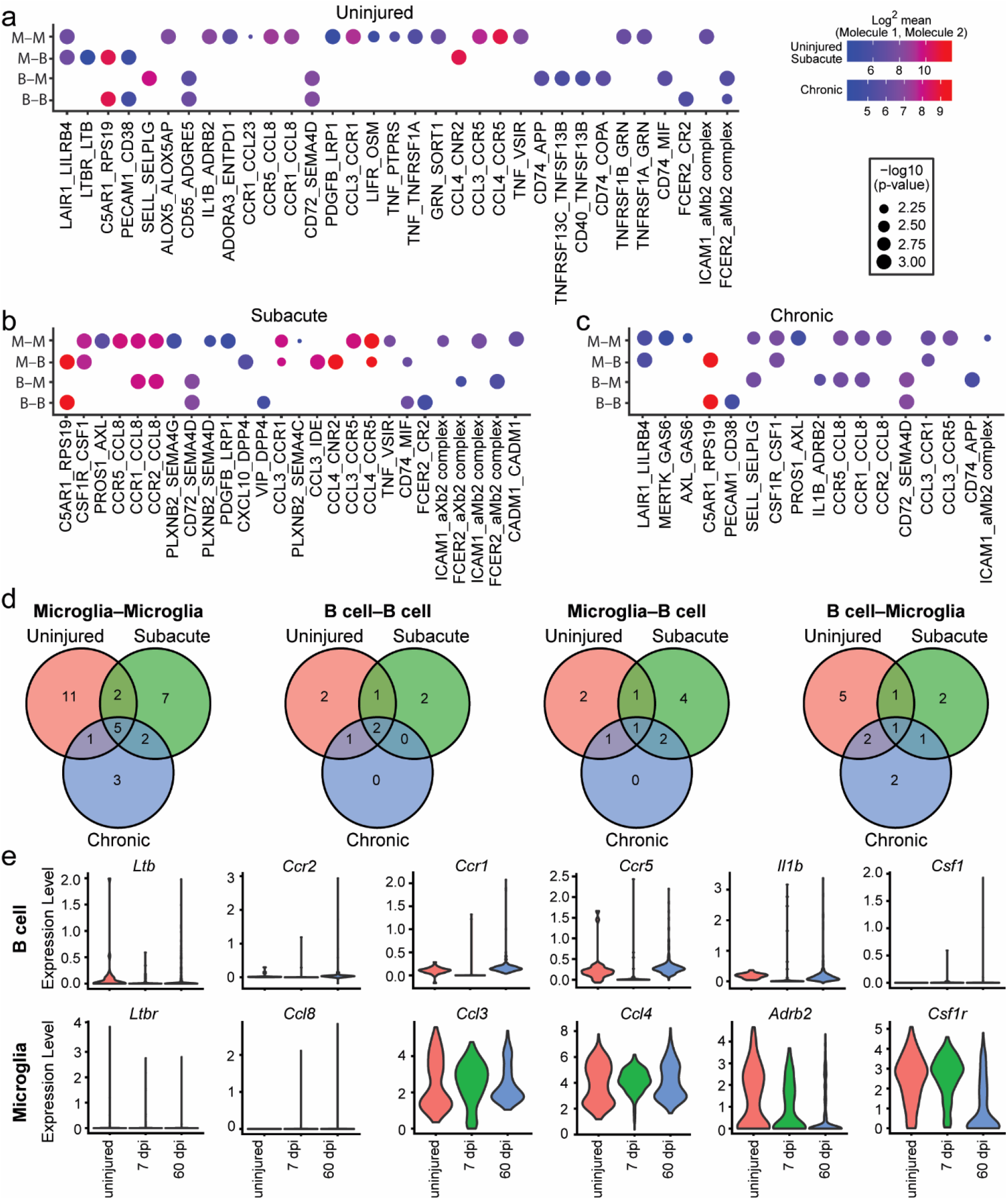
Cell-cell interactions between microglia and B cells in the injured spinal cord. a-c) Microglia (clusters 0,1,5,9) and B cell (clusters 2,3 and 7) gene expression was analyzed at each timepoint (a) uninjured, b) subacute, c) chronic) using CellphoneDB^47^ to determine genes involved in cell-cell interactions, shown as a dot plot. d) Summary of the number of interactions within and across cell types and how they change over time using Venn Diagrams (See **Supplemental Table 5** for lists of these pairings). e) Violin plots of selected genes discussed in text, and their expression in B cells or microglia.

In line with functional identification of “negative regulation of viral process” in microglia from the uninjured spinal cord (**Figure 6c**), one of the seven unique B-M/M-B interactions in uninjured spinal cord was the M-B interaction of lymphotoxin B receptor (*Ltbr*), and *Ltb* (**Figure 7a**). In the lymph node, LTB–LTBR signaling between B cells and macrophages maintains macrophage responsivity to viral infections^48^, and microglial LtBR has roles in de- and remyelination^49^. After SCI, 11 unique interactions between microglia and B cells (M–B/B–M) were found (**Figure 7d)** and notably, most were prevalent in subacute SCI (subacute, n=6; chronic, n=2; subacute and chronic, n=3).

More than half of the interactions identified involved chemokines, potent attractants for peripheral immune cells to injury sites^22, 50^. Microglia expression of the chemokines *Ccl3*, *Ccl4*, *Ccl8* and *Cxcl10* comprised seven of the B-M/M-B interactions identified. In subacute SCI, interactions were found between microglia-expressing *Ccl8* and B cells expressing receptors *Ccr1* or *Ccr2*, and between microglia-expressing *Ccl3* or *Ccl4* and B cells expressing *Ccr1* or *Ccr5* (**Figure 7b**). In chronic SCI, the interactions for *Ccl3*- and *Ccl8*-expressing microglia and *Ccr1*- and *Ccr5*-expressing B cells persisted, and *Ccr1*- and *Ccr5* expression in B cells was augmented (**Figure 7e**). *Ccl4* interactions were not observed in chronic SCI, but a new chemokine-receptor interaction emerged between *Ccl8*-expressing microglia and *Ccr5*-expressing B cells. The functional effects of these chemokines on microglia-B cell interactions after SCI remains to established. However, CCL8 is linked to leukocyte chemotaxis in other inflammatory conditions^51–53^, and SCI- *Ccl3*-/- mice recruit fewer immune cells to the injury site^54^, indicating microglia might attract B cells via CCL8 production; this and other B–M/M–B interactions will be worthwhile investigating in functional studies.

Interestingly, of B cell-related pathways in chronic microglia (**Figure 6c**), only two of the identified interactions were unique to chronic SCI. One was *Ccr5*–*Ccl8*, discussed above, the other was an interaction between *Il1b* in B cells and *Adrb2* (adrenergic receptor β2) in microglia (**Figure 7c**). In dendritic cells, *Adrb2* activation promotes production of anti-inflammatory lymphocytes^55^, and in microglia it is neuroprotective^56–58^. One possibility is that chronic SCI microglia, rather than instructing B cells to invade or cluster as predicted to occur in subacute stages, are exchanging anti-inflammatory signals.

In line with possible protective roles for B cells, in both subacute and chronic SCI, an M-B *Csf1r*–*Csf1* interaction was identified (**Figure 7c**). Colony stimulating factor 1 receptor (CSFR1) signaling contributes to microglial survival^59^; and a study in humans found CSF1 is expressed by activated, but not resting B lymphocytes^60^. This suggests post-SCI, B cells produce CSF1 that promotes microglial health. Hence, the presence of activated B cells could be important for the long-term survival and maintenance of microglia after SCI.

## Discussion

Secondary injury propagated by immune cells leads to incomplete wound healing and chronic inflammation in the spinal cord. By performing scRNA-seq on CD45^+^ cells isolated from the adult mouse spinal cord in the uninjured, subacute, and chronic phases of SCI, we have revealed previously unappreciated diversity and changes in immune cell subtypes over time post-SCI.

Peripheral immune cells increased significantly in the spinal cord following injury, including myeloid-derived-, NK, T, and B cells. Except for myeloid cells, most peripheral cells peaked at 60 dpi, highlighting the importance of studying chronic inflammation in the spinal cord. One drawback of our study is the limited number of cells of the myeloid lineage isolated, possibly because our gating strategy excluded larger, more granular CD45^+^ cells (**Supplemental Figure 8**). Macrophages, neutrophils, and dendritic cells all play crucial roles in modulating the wound environment, and the limited number of these cells precluded deep analysis of genes and pathways expressed, although several studies have begun to elucidate these mechanisms^3, 6^.

At the single-cell level, microglia predominated in the uninjured and subacutely injured spinal cord but were reduced in number chronically. We found in the uninjured state, microglia in the spinal cord were different from those in the brain, and their response following SCI differed from the response of brain microglia to injury or disease. Such differences may help explain reports that similar magnitude injuries induce a less severe immune response in the brain compared to the spinal cord^32^. This contrasts previous work showing similarities between SCI and DAM microglia^4^, though this may be due to injury severity or mouse strain used.

The largest change identified in response to SCI was the ∼6-fold increase in B cell number chronically, which occurred within the injury site, as reported previously^21^. This was accompanied by an increase in B cell diversity: we observed multiple stages of the lineage including pro/pre B and mature B cells. Immature, actively rearranging pro/pre B progenitors were not observed in the uninjured state but appeared in the subacute phase following injury. The presence of proB2 cells could explain how deleterious antibodies against autoantigens are generated following SCI, as this is the stage when B cells are undergoing V(D)J recombination^41^. The expression of MHC-II and changes in Igh expression could participate in the formation of immune complexes linked to complement-mediated cytotoxicity within the chronically injured spinal cord^21^. At all stages, mature B cells were detected with diverse post-class switching immunoglobulins, as well as those expressing MHC-I and -II molecules.

Ectopic clusters of B cells form within the spinal cord in autoimmune disease^61^, and can occur within the injured spinal cord parenchyma^21, 41^ and meninges^43^ and generate cytotoxic antibodies^21, 62^. It is tempting to suggest B cells are recruited during subacute SCI, and respond to injury by setting up ectopic lymphoid follicles that grow locally, producing B cells reacting to the injured environment by producing antibodies and other factors contributing to long-term inflammation^61^. Indeed, the shift from B cells coalescing in ‘ring-like’ structures at 28 dpi (**Supplemental Figure 3c**) to large foci at 42 dpi suggest these clusters grow from B cells initially invading the spinal cord. The presence of B cells in different stages of maturation and the functional identification of IL-23 mediated signaling lends further support for this possibility, as IL-23 drives germinal center formation^45^. In other tissues, ectopic lymphoid follicles resolve after antigen clearance^63^. It would be interesting to examine spinal cords 90 dpi and beyond to determine when, or if, these structures resolve. Persistence of B cells at 60 dpi suggests ongoing B cell stimulation and antigen presentation. Prior attempts to identify the antigens have been unsuccessful, but showed they likely don’t include myelin basic protein^21^. Recently, GlialCAM was linked to multiple sclerosis pathology and the induction of a robust B cell response that aggravates EAE^64^. It would be valuable to identify antigens stimulating B cells after SCI and determine whether GlialCAM is also involved in SCI.

The mechanism leading to B cell invasion into the spinal cord following SCI is currently unknown, although we show chemokine signaling may be involved. Intriguingly, other studies have shown lymphopoiesis is impaired in the bone marrow following SCI and B cells fail to mobilize^65^, suggesting B cells observed in the injured spinal cord might have a different origin. Indeed, recent studies suggest the meninges, cerebrospinal fluid, and vertebral bone may be a source of B cells entering the injured CNS^66–69^. Our study showed early B cells in the spinal cord parenchyma are similar to those found in the spinal cord meninges^43^, underscoring this as a possible source. Identification of the mechanisms whereby B cells enter the spinal cord, and processes associated with B cell development and activation within the injured spinal cord should help define their role in cell toxicity and allow more nuanced targeting of B cells following injury.

We uncovered several interactions between immune cells, specifically, between microglia, neutrophils, and B cells that may contribute to the retention and accumulation of B cells at the injury site. From the combined functional and CellPhoneDB changes identified, we hypothesize altered microglia-neutrophil interactions signal the early invasion of neutrophils^70^ that in turn initiates subacute entry of B cells at the site of SCI^71^, which is augmented chronically by signals generated by reactive astrocytes and microglia. Identifying the signals initiating these changes is needed to refine and test this hypothesis and to develop targeted interventions. This dataset could be used a reference for further investigations into resident-peripheral immune interactions in acute and chronic SCI, including those of T cells which were not evaluated in this study. Here we identify several chemokines, important cell attractants, as possible receptor-ligand interactors between microglia and B cells. Future work should focus on identifying interactors between B cells and other immune cells and glia, and test the cell-specific expression and functional roles of chemokines identified here on the B cell retention and function at acute and chronic stages of SCI.

## Online Methods

### Animals

The care and use of animals were approved by UAlbany IACUC and overseen by the Animal Resources Facility at the University at Albany East Campus Animal facility.

### Spinal cord injury

Adult (10-12 wk. old) female Swiss Webster mice were housed in a pathogen-free barrier facility and maintained on a 12-hour dark/light cycle with food and water provided ad libitum. First, mice were deeply anesthetized by inhalation of isoflurane vapor (3%). The contusion SCI surgery was performed as described^72^ using the NYU Impactor. Spinal cords were exposed by laminectomy at T9-10 levels and contused by a 10 g rod dropped from 6.25 mm. Immediately after the injury, muscle was approximated using interrupted chromic gut sutures, skin was closed with interrupted chromic gut sutures, and lidocaine gel was applied. After surgery, mice were maintained on heating pads, closely observed until fully awake and then returned to their home cages. Subcutaneous Buprenex (0.2 mg/kg) was injected twice daily for three days for postoperative pain control. Bladders of injured mice were expressed twice daily. At 3, 7 and 60 days after SCI, animals were sacrificed by decapitation under deep anesthesia with 80 mg/kg pentobarbital.

### Flow cytometry

Following sacrifice, animals from all groups (n=6/ timepoint) had a 3 mm section of spinal cord dissected out centered at T9. Meninges were removed from the spinal cord segments under a dissecting scope. The segments were pooled and minced in ice-cold Hank’s Balanced Salt Solution (HBSS). Following enzymatic dissociation with trypsin (0.25mg/mL in DMEM) at 37° for 20 min, the cells were triturated with a Pasteur pipette to further dissociate the cells and then passed through a 40 μm cell strainer to reduce myelin debris. Trypsin was inactivated by the addition of DMEM + 10% Fetal Bovine Serum (FBS). Cells were then centrifuged and washed twice with fluorescence activated cell sorting (FACS) buffer (phosphate-buffered saline (PBS), 2% bovine calf serum). This was followed by a 30 min incubation with antibodies against CD45 (FITC, BD Biosciences #553079) diluted 1:20 in FACS buffer, shielded from external light. The cells were then washed twice with PBS. Flow cytometric analysis was performed using a BD FacsAria Flow Cytometer/Cell Sorter. Sorting gates were determined based on unstained, uninjured control animals. CD45^+^ Hi and Lo populations were selected and both groups collected into a 1.5 ml tube containing DMEM.

Gating selection for the CD45 positive immune cells of the spinal cord is illustrated in **Supplemental Figure 7**. Doublet discrimination was performed twice to remove as many as possible from the samples.

### iCell8 preparation and dispense

Cells were prepared according to the iCell8 protocol prior to single-cell dispense. Briefly, the total cell number obtained from FACS was used to calculate a dilution to dispense 1400 cells per well in a 384 well plate. Cells were stained with NucBlue (identifies cell nuclei during the imaging step) at a concentration of 2:25 and Propidium Iodide (to identify dead cells) at a concentration of 1:25 for 20 min at 37°C (ThermoFisher Cell Viability Imaging Kit #R37610). Cells were centrifuged for 5 mins at 300 g, and supernatant was discarded. Cells were resuspended in 80 µL/well dPBS. 1 µL/well of RNase (New England Biolabs # M0314L) was added, plus 1 µL/well iCell8 100X diluent. 80 µL of resuspended cells was added to each of the designated sample wells of the iCell8 384-well plate. The 384 well plate and an iCell8 chip were loaded into the iCell8 machine, and the standard dispense program was run to dispense 50 nL of sample into each well of the chip (total 5184 wells), in addition to fiducials (dye used to center the well for the imaging step for cell selection) and RNA controls (Takara #636643).

### Chip imaging and cell selection

After dispensing, the chip was blotted with filter paper and sealed with imaging film. The chip was centrifuged at 300 g for 5 min and set on the microscope. Fiducials were confirmed and images acquired via iCell8 Image software. Once images were acquired, the chip was placed in a chip holder and frozen at -80°. Single, live cells were selected based on imaging for NucBlue/ propidium iodide stain using iCell8 CellSelect software (**Figure 1c**).

### Single-cell RNA library preparation and sequencing

From uninjured control, 3 dpi and 7 dpi, 500 spinal cord immune cells were selected from each chip as single cells. Five K562 cells (control cells provided with the system), five negative control wells (no RNA, but all iCell8 buffers), and five positive control wells (12 pg RNA) were also identified. For 60 dpi, 1000 spinal cord immune cells were isolated and dispensed in a similar manner.

Libraries were prepared according to the iCell8 single-cell SMART-seq 3’ DE library preparation kits (Takara #635040) and were sequenced (150 cycles) on the NextSeq500 platform (Illumina) lane (4x10^6^ potential total reads).

A Bioanalyzer was used to assess the quality of the resulting cDNA and to monitor the size distribution of the transcripts. From our samples we obtained approximately 7-8 ng of cDNA with an average of 1650 base pair fragments, indicating good quality sampling of the immune cell RNA from the uninjured and injured spinal cords, enabling us to proceed to sequencing of the cDNA.

### Analysis of single-sell RNA sequencing data

Data was initially processed using Bcl2fastq (Illumina) to convert raw BCL files to fastq. Fastq files were split based on single-cell barcodes and iCell8 well list files using in house java and R code, which is available on github (https://github.com/neural-stem-cell-institute/sc-pipeline). Reads were mapped to mouse genome version mm10 using STAR aligner (v2.5)^73^. Aligned reads were sorted and aggregated using SAMtools and raw counts determined using Bioconductor packages in R^74, 75^ .

Following the well documented workflows of the Seurat3 package^76^ we normalized, scaled, transformed, integrated, and performed cell-cycle correction on the raw read data. Subsequently, the Seurat3 library was used to perform dimensional reduction, neighbor-finding, clustering and then to identify marker genes for the resulting clusters. Markers were computed for each cluster relative to the complete dataset. Pairwise markers were computed to compare clusters determined to be comprised of subtypes of the same general cell identity. The SingleR and hypeR annotation and enrichment tools were used to determine functional enrichment of marker genes and top-expressing genes by cluster and by time point^29, 77^.

#### Gene ontology enrichment analysis

Differentially expressed genes and top expressing genes from time points and from specific cell-type clusters were used to determine enriched gene ontology categories. These gene lists were processed using Gprofiler2 and Revigo^78, 79^ to calculate enrichments. Enrichments and plots generated for **Figure 6 a,b** and **Supplemental Figure 6** were done using the Screp function (https://github.com/neural-stem-cell-institute/screp).

#### Integrated analysis of microglia with data from Hammond, et al^35^

Whole brain control and lysolecithin stimulated data sets published by Hammond et al.^35^ were retrieved from the Gene Expression Omnibus (accession: GSE121654). The Seurat library was used to integrate these data, along with the data from our study, using the SCTransform library (which performs normalization, variance stabilization, and feature selection based on a UMI-based gene expression matrix). After this normalization, SCT dimensional reduction and clustering and cluster marker identification was performed using the well documented methods RunUMAP, FindNeighbors, FindClusters and FindMarkers from the Seurat library.

#### Integrated analysis of B cell subtypes with Jensen data

Cells from this study that we identified as B cells were analyzed independently from the remainder of our data set. The SingleR Bioconductor package^29^ was used to create a reference dataset from Gene Expression Omnibus (accession: GSE74290). The SingleR methods were then applied following documented workflows to label B cells from our study relative to the reference dataset we constructed. To highlight the most stable labels, a range of score cutoffs were evaluated and a score cutoff of 0.125 was selected to indicate the B cell subtype labels generated in the reference.

#### Integrated analysis of B cell subtypes with Cohen data

Data published in the gene expression omnibus by Cohen et. al^43^ (accession: GSE160193) provided another reference dataset for the determination of potential labels for the B cells collected in this study. Sample GSM4862209 was excluded as its smaller scale severely limited the number of integration anchors that could be calculated during this integration. Cells were integrated into a single combined Seurat object along with all the cells collected in our study. The Seurat library was used to integrate this data following a process of SCT normalization followed by dimensional reduction and clustering. The relative location of labeled cells from Cohen^43^ (“B” and “pre B”) and the B cells from this study (**Supplemental Figure 6**) were observed via UMAP to infer likely cell type convergence.

#### Integrated analysis of B cell subtypes with TabulaMuris data

Immune-relevant data subsets were collected from the *TabulaMuris* Consortium data^80^. This included both droplet and FACS sorted cells from spleen, marrow, brain, fat, limb, liver, and lung. The data was normalized and combined into one Seurat object using a process of finding the top 2000 variable features and 20 integration anchors. Smaller data sets from *TabulaMuris* cohort constrained the total number of reference anchors that could be utilized in this calculation. Seurat methods FindTransferAnchors and TransferData were used to find anchors and transfer labels was then used to predict labels for our B cells.

### Data and code availability

Raw and processed data is available for download from the Gene Expression Omnibus (GEO) (Currently under submission). Processed RData files, suitable for exploration within the R programming environment, along with sample interaction code, as well as code for the above analyses, can be obtained from GitHub via our institution website at http://neuralsci.org/computing.

### Spinal Cord Staining

Spinal cord tissue sections were kindly provided by Drs. John Gensel and Andrew Stewart from spinal cords harvested and sectioned from a previous study^81^. Briefly, male C57/Bl6 were given a 60 kDyn SCI under ketamine (100.0 mg/kg) and xylazine (10.0 mg/kg) anesthesia using the Infinite Horizons Impactor (Precision Systems Instrumentation, LLC, Fairfax Station, VA). Mice were anesthetized using an overdose of ketamine and xylazine and perfused using cardiac puncture with 0.1 M PBS, then with 4% formaldehyde in PBS. Spinal cords were extracted, then post-fixed for 2 h at room temperature and washed in 0.2 M phosphate buffer overnight. Cords were transferred to a 30% sucrose solution for 1 week before blocking in optimal cutting temperature compound (OCT; Thomas Scientific; Swedesboro, NJ). 10.0 μm thick sections were cut in the coronal plane between −18°C and -20°C on a cryostat.

Slides with coronal spinal cord tissue sections were removed from -20°C storage and dried at room temperature for 1 hr. Tissue sections were then permeabilized for 1 hour and blocked in PBS with 5% normal goat serum, 3% BSA, and 0.03% Triton-X 100. The background was quenched using TrueBlack (BioTium # 23007) according to manufacturer instructions. B cells were stained for CD45RA (BioRad # MCA1258GT) diluted 1:100 and neurons for Tuj1 (BioLegend #801202) 1:500 in the same blocking solution and incubated overnight at 4°C, followed by three washes with PBS. Secondary antibody (Jackson Labs #712-545-153 (B cells), Invitrogen #A21135 (neurons)), was added at 1:500 in blocking solution for two hours at room temperature followed by three washes with PBS. DAPI (Invitrogen #D1306) was added 1:1000 in water for 5 mins, followed by two washes in PBS. Tissue was mounted in Fluoromount (Sigma #F4680). Tissue was then imaged using a Zeiss LSM 780.

## Acknowledgements

We thank the U. Albany Center of Functional Genomics, including Dr. Sridar Chittur and Marcy Keuntzel for performing RNA sequencing. We are grateful to Andrew Stewart and John Gensel for their generous gift of tissue sections used in Figure 5. We thank the members of the Temple lab for input on the manuscript and intellectual discussion, Nathan Boles for bioinformatic guidance for E.S.F., and Jonathan Adamec for assistance with histological quantification of B cells. Supported by the Spinal Cord Injury Trust Fund through New York State Department of Health Contract #C32244GG. Opinions expressed here are solely those of the author and do not necessarily reflect those of the Spinal Cord Injury Research Board, the New York State Department of Health, or the State of New York.

## Author Contributions

S.T. and T.R.K. conceptualized the study, M.A. and N.L. performed the SCI and cell isolation, M.A and S.L. performed and assisted with the FACS sorting and iCell8, respectively. M.A. generated the library. T.R.K., E.S.F and F.F. performed the bioinformatics. E.S.F, C.E.H., T.R.K, N.L, S.T. analyzed and visualized the data and interpreted the results. E.S.F and C.E.H performed the histological B cell analysis and imaging. E.S.F, C.E.H., T.R.K. generated the figures. E.S.F generated the first draft of the manuscript, which C.E.H., and S.T. edited. T.R.K, M.A., and N.L. provided critical feedback on the manuscript.

## Supplemental Figures

**Supplemental Figure 1.**
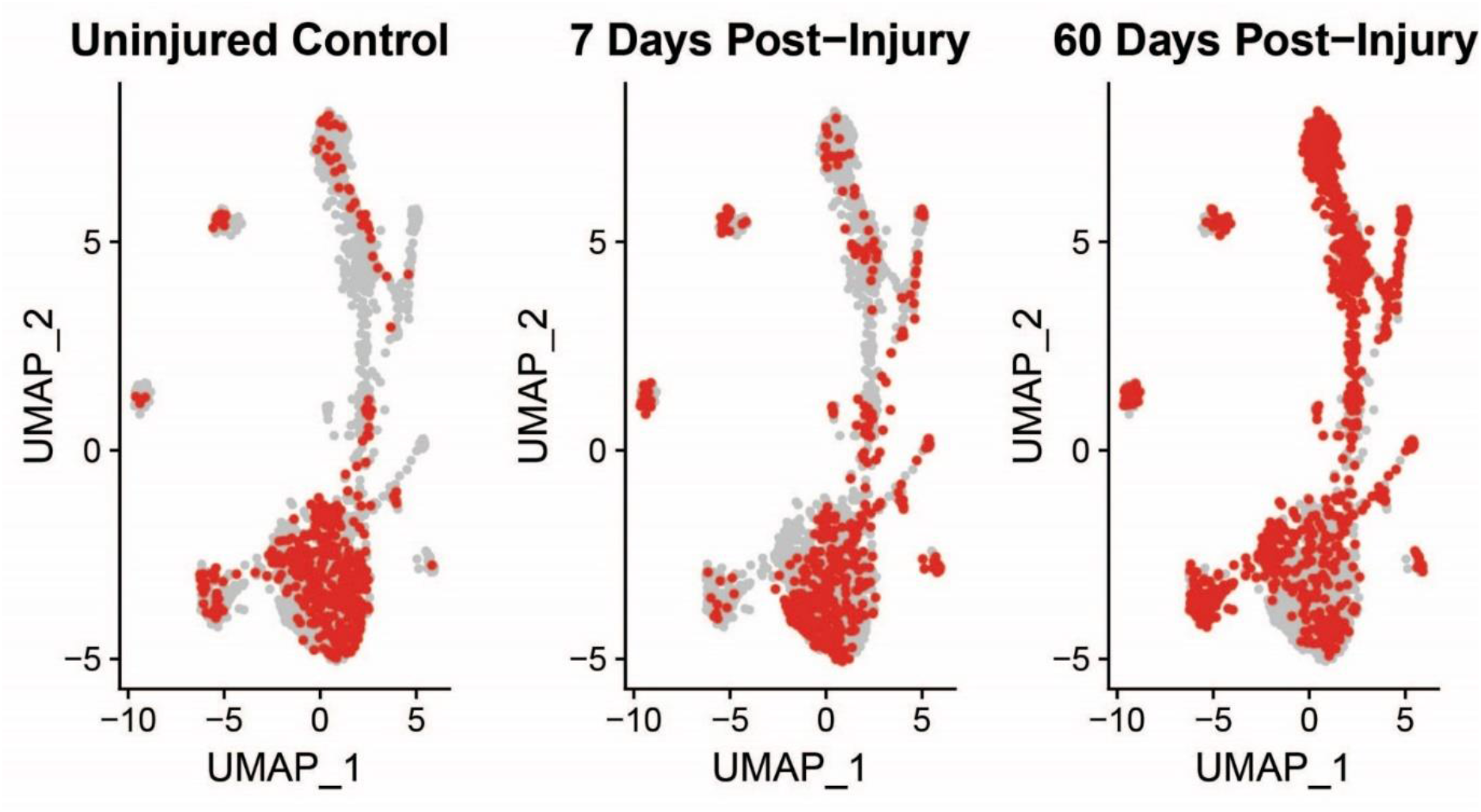
Representation of cells across timepoints on the UMAP after SCTransform. All cells collected at the control, subacute (3 and 7 dpi), and chronic (60 dpi) mapped onto the UMAP of all cells together.

**Supplemental Figure 2.**
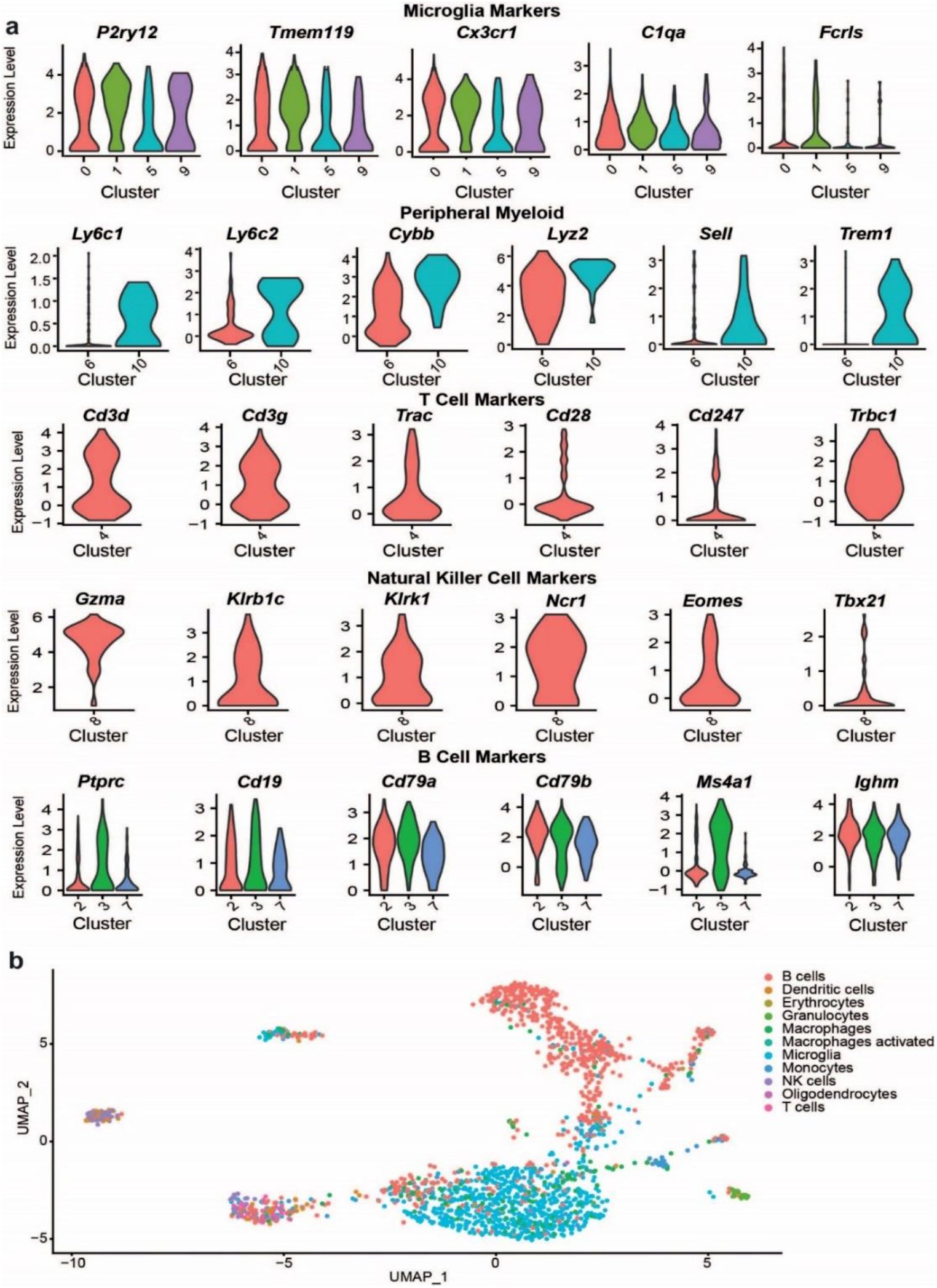
Confirmation of cell identity and markers of each cluster. a) For each cell type identified (microglia, macrophages/monocytes, T-cells, NK Cells, and B cells), canonical markers were examined via violin plots to confirm their classifications. b) Cells isolated from the spinal cord were computationally compared to immune cells in the ImmGen database, confirming that most cells isolated were microglia or B cells.

**Supplemental Figure 3.**
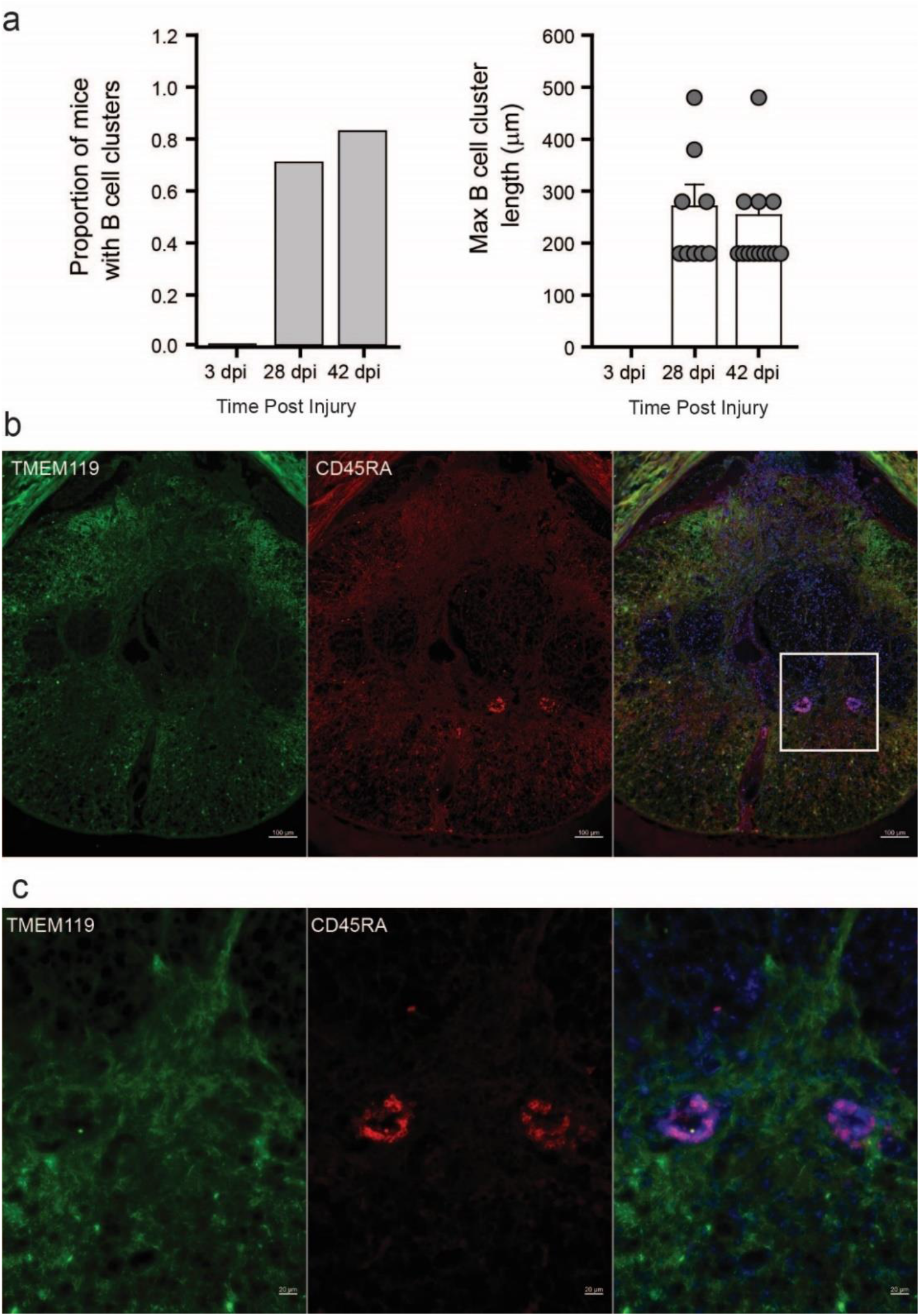
B cell cluster analysis in tissue sections. a) Quantification of proportion of mice analyzed with B cell clusters present anywhere within the lesion site. b) Quantification of the maximum length of B cell clusters present within the tissue. c) Immunohistochemical staining of spinal cord sections for TMEM119 (microglia) and CD45RA (B cells) from an animal 28 dpi.

**Supplemental Figure 4.**
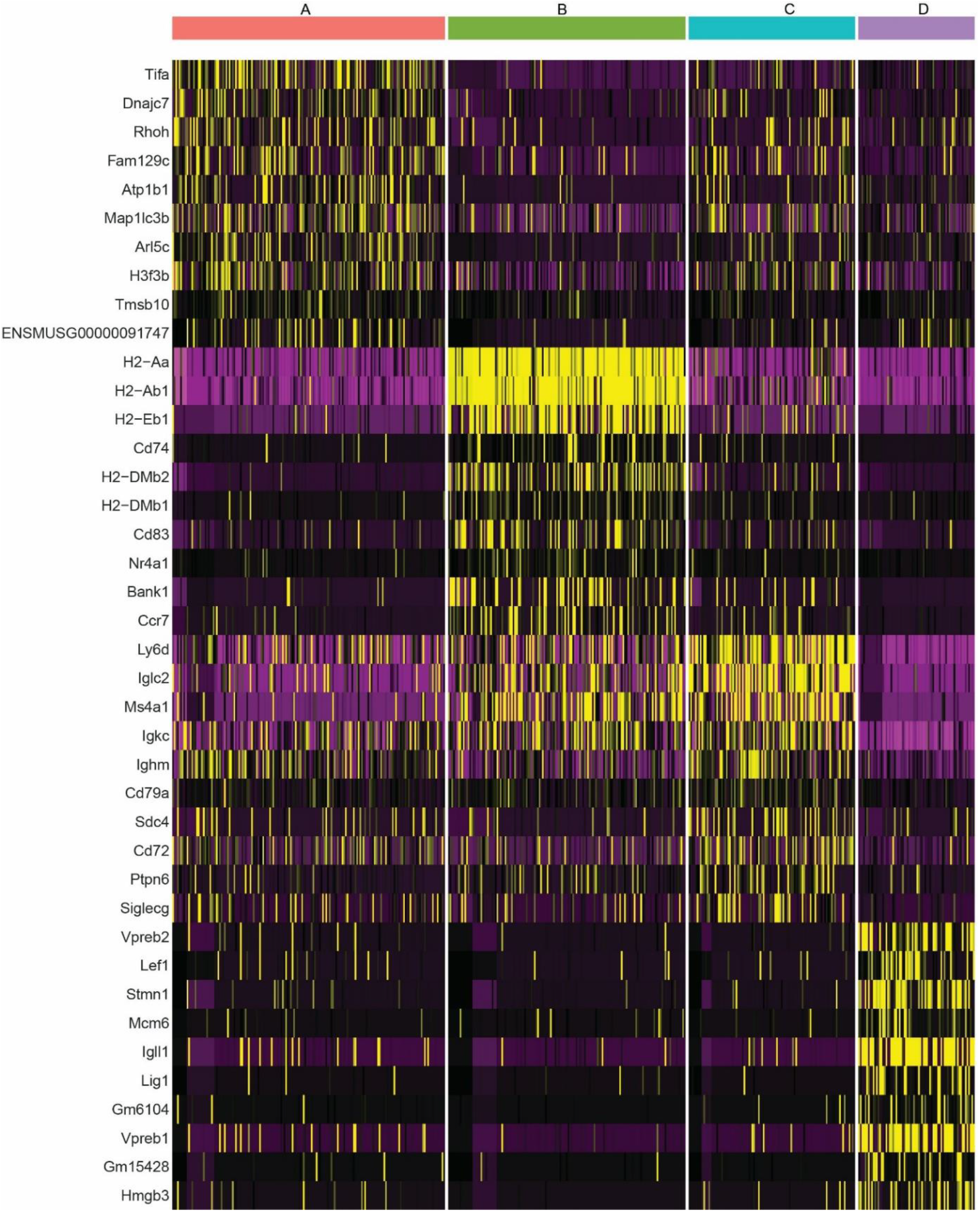
Heatmap of marker genes for B cell clusters identified in Figure 5.

**Supplemental Figure 5.**
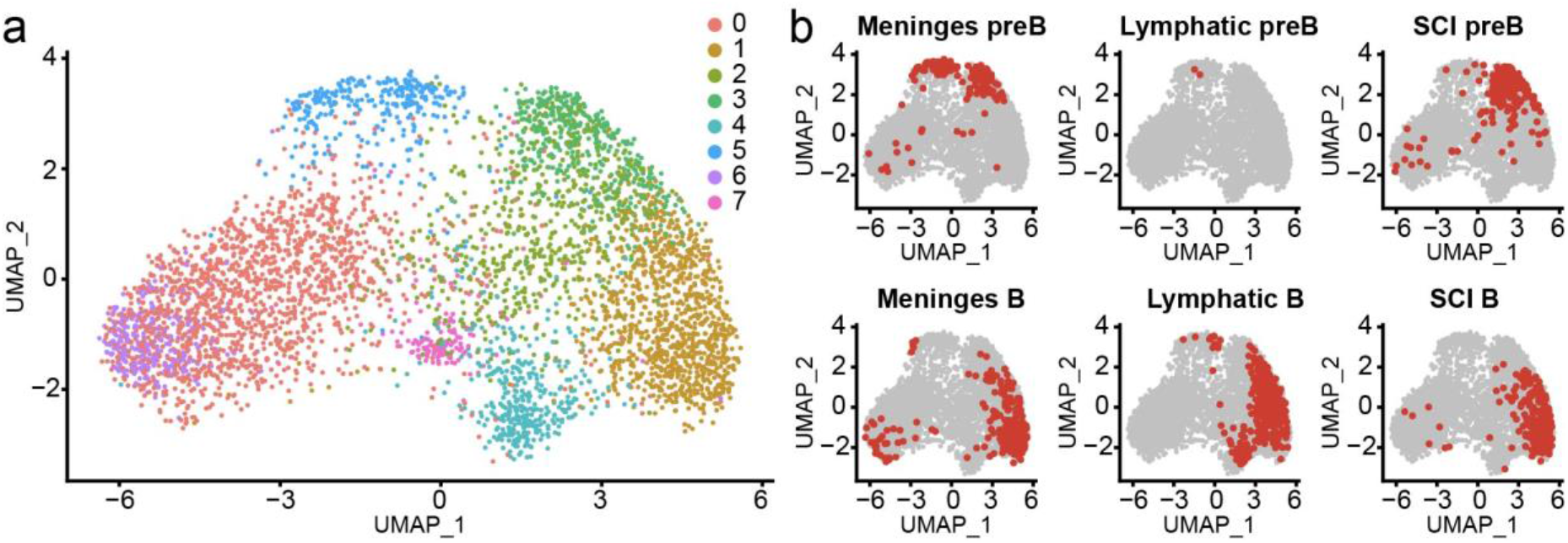
Analysis of B cells isolated from SCI combined with those isolated from the meninges. a) UMAP of all immune cells isolated from the spinal cord in our study (uninjured and injured) combined with the data from Cohen et al. b) Mapping of subpopulations of B cells (pre B and mature B) identified in meninges, lymph nodes (lymphatic), and from SCI.

**Supplemental Figure 6.**
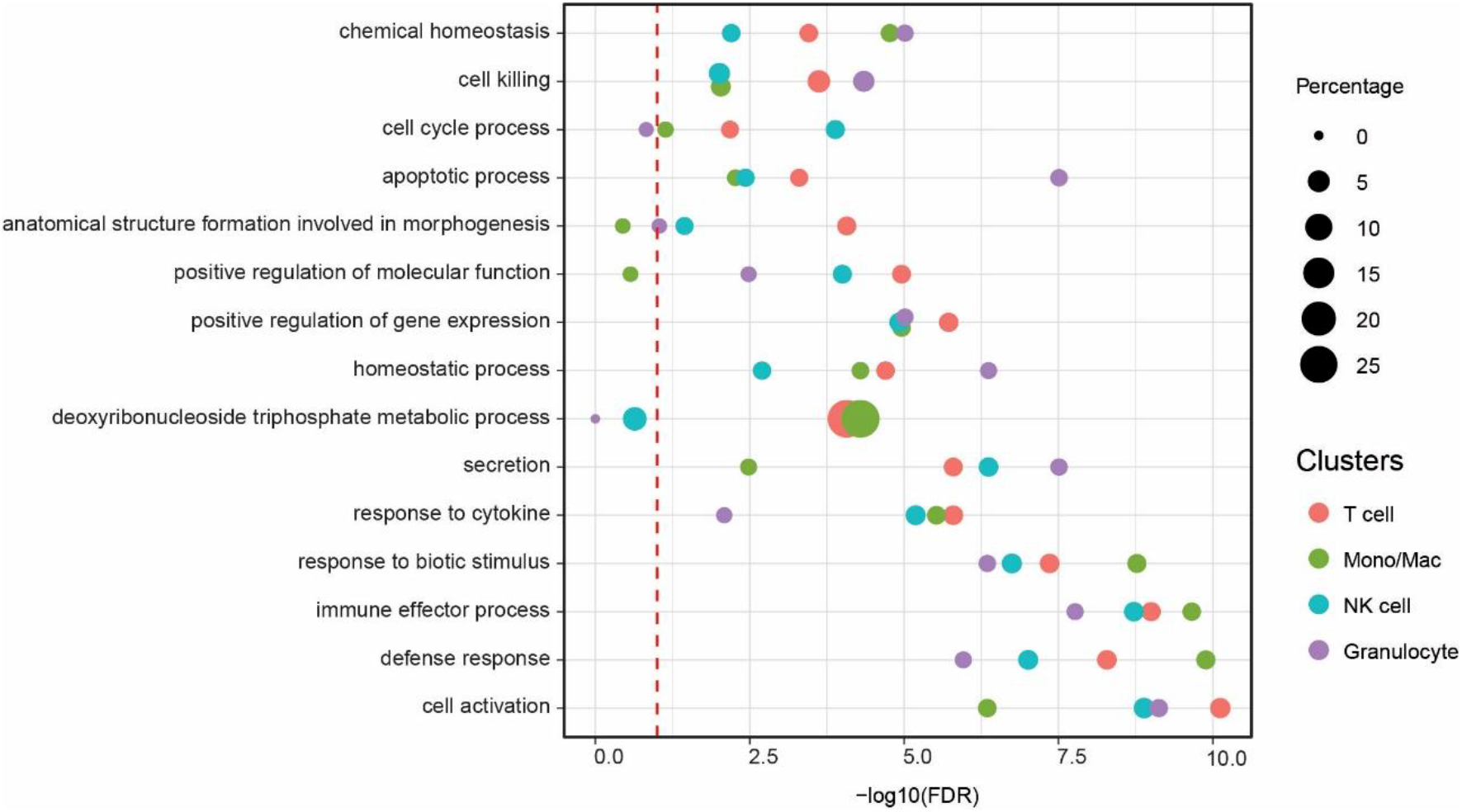
GO terms enriched in clusters for T cells, Monocytes/macrophages, NK cells and Granulocytes. Wilcoxon rank sum test was used to identify markers for each cell type. Hierarchical groupings were performed as in Figure 4, and a dot plot was generated. The size of the dot indicates the percentage of gene overlap, and the x-axis is FDR. Red dashed line indicates FDR=0.1. See **Supplemental Table 6** for additional enrichments and genes associated with the enrichments.

**Supplemental Figure 7.**
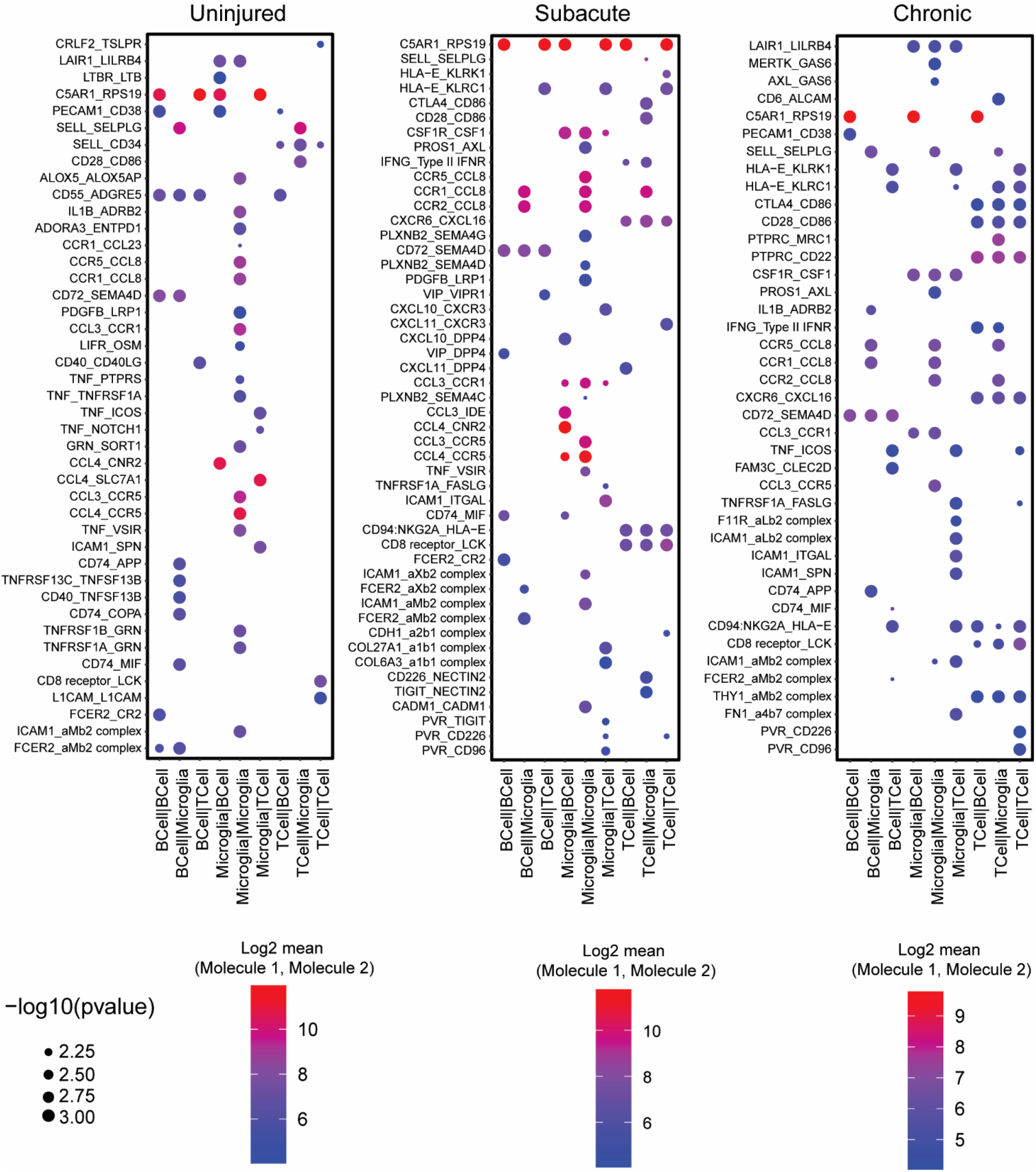
CellPhoneDB interactions between microglia, B cells, and T cells. CellphoneDB was used to determine receptor ligand pairs between microglia, B cells and T cells at the uninjured, acute, and chronic phase of injury. Results are plotted for interactions which the p<0.01.

**Supplemental Figure 8.**
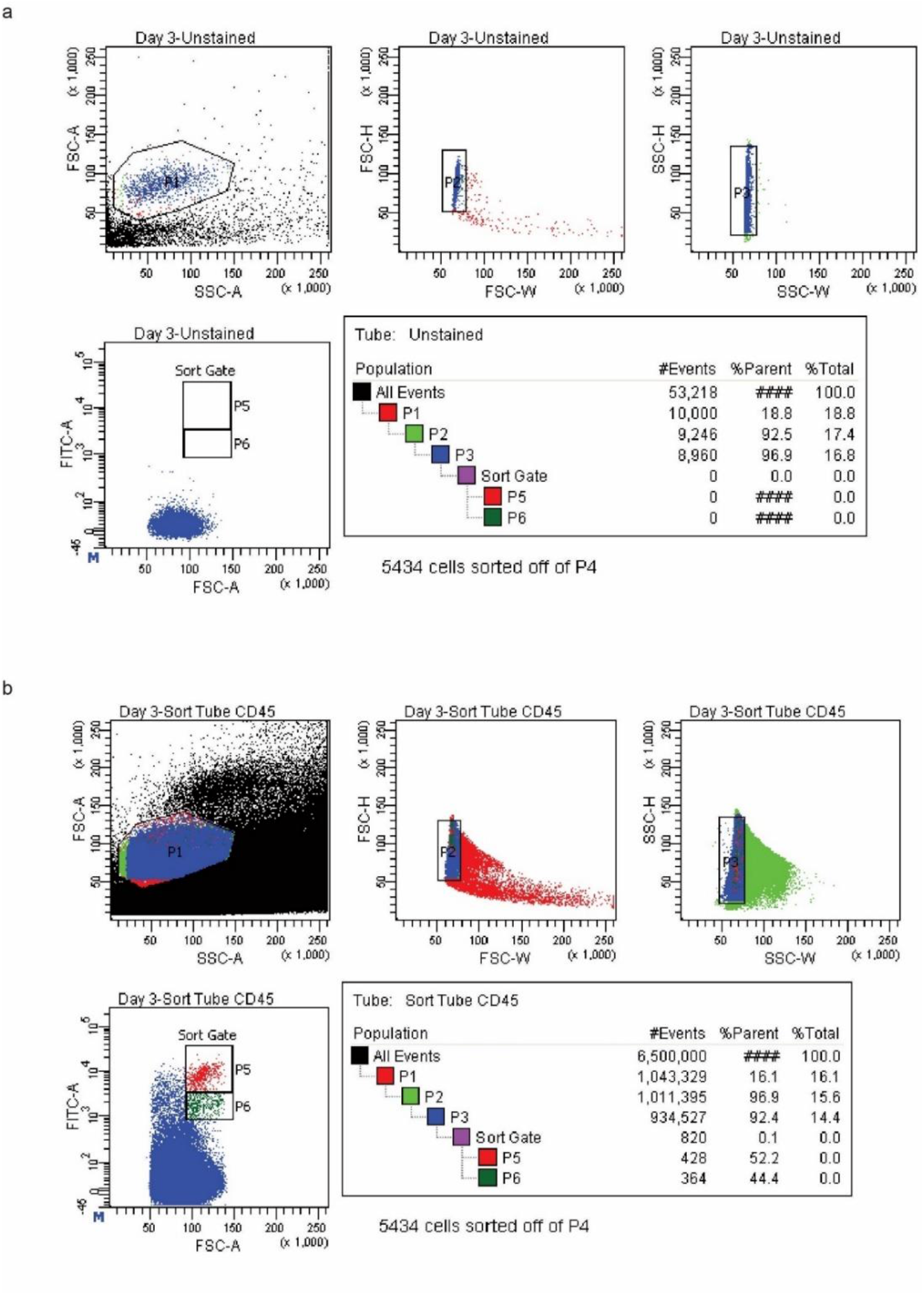
Gating and flow sorting parameters for isolating single CD45^+^ cells. a) Unstained spinal cords were used to set the gates and to establish single cell isolation. P1 identifies the region of live cells. P2 and P3 are forward and side scatter gates to identify single cells. P5 and P6 are the CD45^+^ cells which were selected for FACS. b) The same schematic with the 3 dpi sample to show isolation of CD45^+^ cells.

